# Neglected Quaternary legacy on biodiversity in the Mountains of Southwest China

**DOI:** 10.1101/2020.02.28.969089

**Authors:** Tao Wan, Huateng Huang, Jamie R. Oaks, Xuelong Jiang, L. Lacey Knowles

**Affiliations:** College of Life Sciences, Shaanxi Normal University, Xi’an, Shaanxi 710119, China; Mammal Ecology and Evolution, State Key Laboratory of Genetic Resources and Evolution, Kunming Institute of Zoology, Chinese Academy of Sciences, Kunming, Yunnan 650223, China; Department of Ecology and Evolutionary Biology, University of Michigan, Ann Arbor, MI 48109, USA; Department of Biological Sciences & Museum of Natural History, Auburn University, Auburn, AL 36849, USA

**Keywords:** comparative phylogeography, Mountains of Southwest China, diversification, riverine barrier, species pump, full-likelihood Bayesian computation

## Abstract

Mountains of Southwest China (MSWC) is a biodiversity hotspot with a very unique and highly complex terrain. However, with the majority of studies focusing on the biogeographic consequences of massive mountain building, the Quaternary legacy of biodiversity for the MSWC has long been overlooked. Here, we took a comparative phylogeography approach to examine factors that shaped community-wide diversification. With data from 30 vertebrate species, the results reveal spatially concordant genetic structure, with temporally clustered divergence events during severe glacial cycles, indicating the importance of riverine barriers in the phylogeographic history of the vertebrate community. We conclude that the repeated glacial cycles are associated with temporal synchrony of divergence patterns that are themselves structured by the heterogeneity of the montane landscape has of the MSWC. This orderly process of diversifications has profound implications for conservation by highlighting the relative independence of different geographic areas in which communities have responded similarly to climate changes and calls for further comparative phylogeographic investigations to reveal the extent to which these findings might apply more broadly to other taxa in this biodiversity hotspot.

## Introduction

Although the uplift of mountains and the associated impact on diversification is widely recognized, including for the Mountains of Southwest China (MSWC), the processes associated with this complex topography and associated barriers as recent drivers of divergence remain relatively less studied. This is especially the case for the MSWC, one of the most topographically complex and species-rich biodiversity hotspots (1). In the past two decades, researchers conducted hundreds of biogeographic and phylogeographic studies in MSWC to study the vicariance events associated with the uplifting of Tibetan Plateau (2), such as the separation between Asian catfish genera (3), and between Senecioneae genera (4). Yet, many cryptic species and recent speciation events have been discovered and documented in this region (5-8), with divergence dates that clearly occur after major orogenesis events in this region (5, 8), suggesting recent divergence may play a role in generating some of the biodiversity in this region, and that mechanisms other than geographic events in the distant past should be considered.

We currently have limited information on recent diversification events and the mechanisms— those responsible for population differentiations and cryptic speciation—in the MSWC. On this temporal scale, these events typically are concentrated in the Pleistocene (9). Spatially, species often exhibit discrete genetic patterns, including those that correspond to geographic or ecological barriers to dispersal, such as arid-hot valleys, basins, or icy mountain tops (10). However, the geographic scope of such studies is fairly limited. For example, most studies only recognize the whole MSWC region as an east-west dispersal barrier. Yet, the MSWC is characterized by a complex of extremely rugged terrain that is divided by three major rivers (the Salween, Mekong, and Yangtze) (Figure 1), forming deep-cutting canyons that are at least 2,000 m lower than the mountain ridges on the two sides. However, a pronounce knowledge gap is associated with the divergence processes at this finer geographic scale. For example, do populations of different species have concordant spatial structure? And if so, is there evidence of synchronous divergence at this finer temporal scale (i.e., recent history)? To what extent are community spatiotemporal divergence patterns similar, as opposed to being associated with species-specific characteristics (e.g., habitat preference)? Do the rivers have similar effects on divergence, or does the effect of the river on community divergence vary (i.e., river-specific effects on species divergence)? It is these questions that motivate our research, and the answers of which, are arguably an important component to understanding the accumulation of species in the MSWC, a biodiversity hotspot.

**Figure 1.**
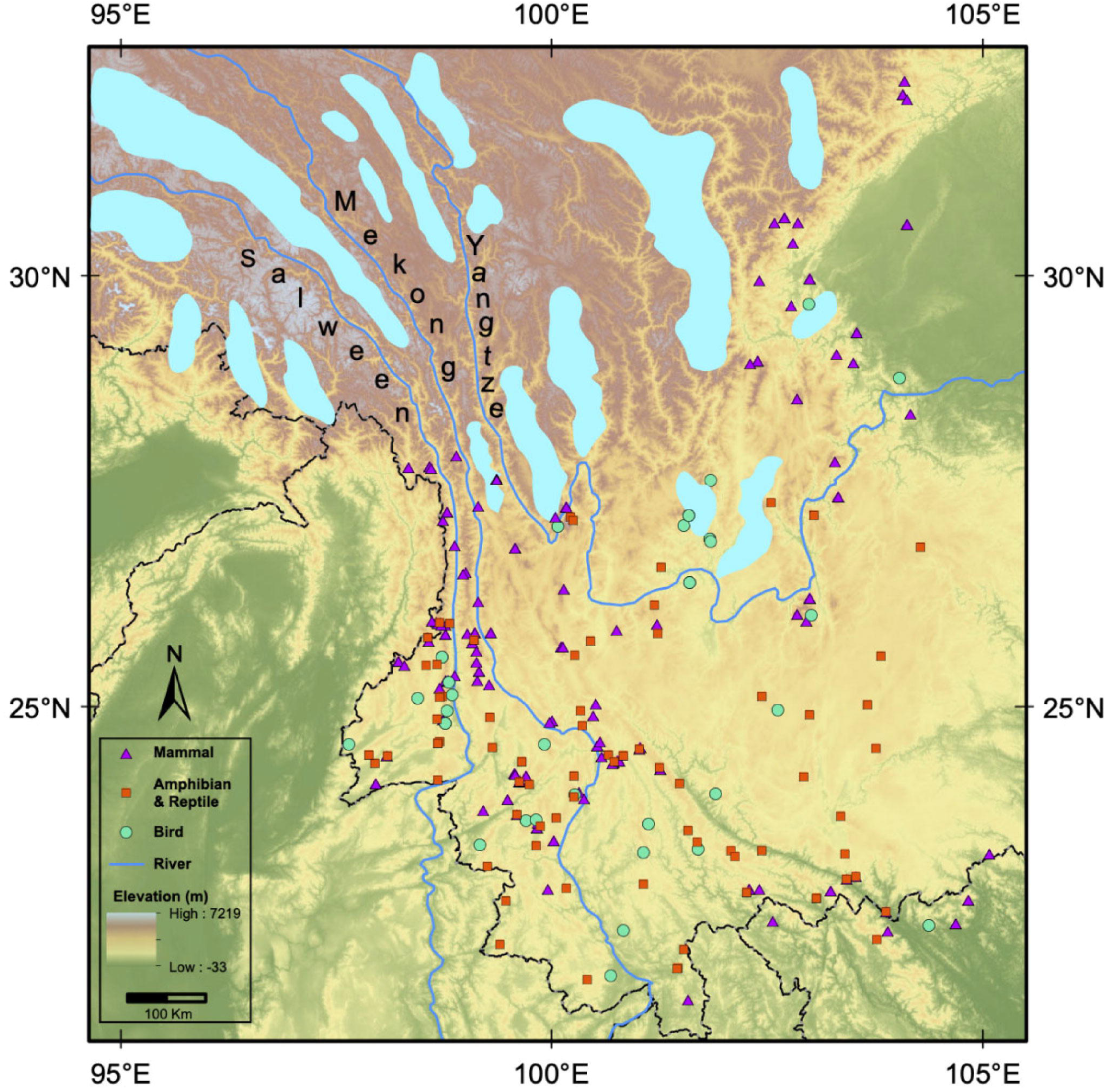
Map of the main rivers and topography of the MSWC. Colored symbols represent the geographic location of the samples; purple triangle, orange square, and cyan circle denote samples of mammals, amphibians and reptiles, birds, respectively. Cyan patches show the maximum extent of glaciation during the Last Glacial, according to Owen and Dortch (2014).

Here, we adopted a community-wide approach to decipher the phylogeographic pattern across multiple co-distributed species in MSWC. The concordance of spatial structures and the synchronization in divergence time in a biological community are used to test hypotheses about mechanisms that generated biodiversity (11-14). Specifically, we analyze data sets generated a new for some mammal species (Table S1 in Supplementary file 1) along with data from previous phylogeographic studies conducted in MSWC (Table S2 in Supplementary file 1) to increase the representation of taxa for tests of concordance of species phylogeographic structure using the full-likelihood Bayesian computation method (FBC; given differences in performance, as well as more general theoretical arguments, we focus exclusively on tests based on FBC, as opposed to relying on approximations in hierarchical approximate Bayesian computation, hABC) (15). For all non-flying terrestrial animals in this study, genetic data reveals that at least one of the rivers functioned as dispersal barriers, with many species exhibiting spatially concordant phylogeographic structure. Hence, we more specifically tested three hypotheses with different levels of temporal synchronization among spatially concordant species: asynchronous hypothesis (H1) in which taxa have independent divergence histories; synchronous hypothesis (H2) in which all taxa “share” one co-divergent event; and partially synchronous hypothesis (H3) in which taxa grouped into more than one co-divergence events (Figure 2). With this set of hypotheses our work aims to address (i) whether there are prevailing phylogeographic histories among co-distributed species in MSWC, and (ii) what factors contribute to community-wide phylogeographic diversity in particular, in the MSWC.

**Figure 2.**
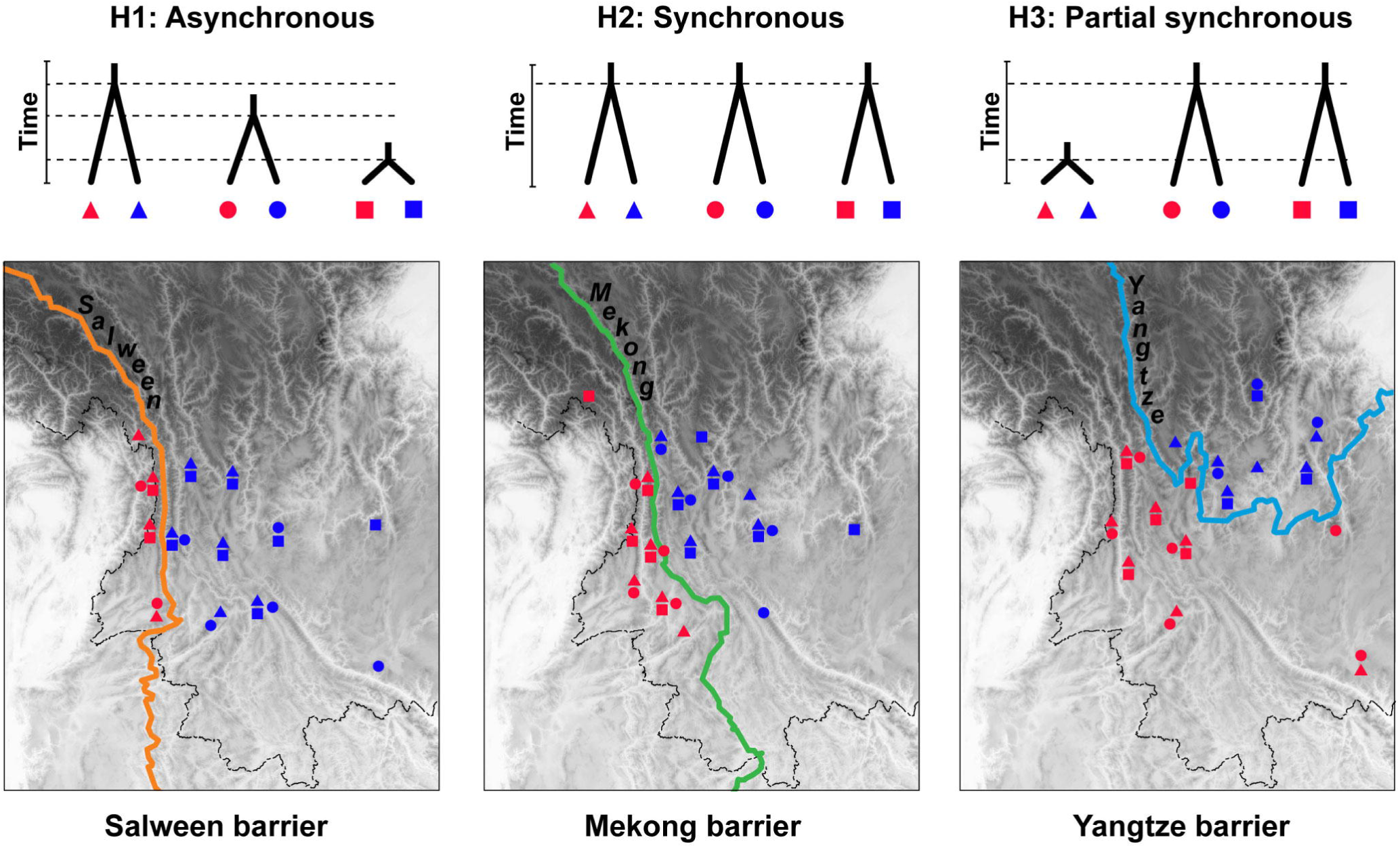
Testing comparative phylogeographic hypotheses among species with spatially concordant phylogeographic structure. The pairs of clades with a phylogeographic break at each river (the Salween, Mekong, and Yangtze) were identified, and hypotheses H1–H3 were tested using the full-likelihood Bayesian computation method. Different symbol shape represents different species; the within-species clades bounded by rivers are represented as contrasting colors.

## Results

### Data acquisition

We took the *sensu lato* concept of the MSWC, which is geographically broader than the biodiversity hotspot MSWC (10). In addition to the sequence data generated specifically for this study, we surveyed the Web of Science using the keyword ‘phylogeog*’ and ‘China,’ keeping only studies with representative sampling across at one of the four focal areas (Figure 1), and for which, there was georeferenced genetic data. Together with the sequence data we generated for six mammal species, the compiled dataset consisted of 2278 mitochondrial sequences from 30 terrestrial species (14 small mammals, 10 birds, and 6 amphibians or reptiles; Table S2). All these species are typical members of MSWC animal community.

### Examining the spatial phylogeographic structure

We used the mitochondrial gene tree estimated by BEAST v2 (16) to examine whether river barriers correspond to monophyletic clades on the gene tree (Figure S1–S3, Supplementary file 2), or were delimited using GMYC (Table S1) (17). Rivers correspond to phylogeographic breaks in all examined mammal species – that is, few clades were distributed across a river (see details in Supplementary Fig S1–S3). Depending on the whole species range, the number of phylogeographic divisions ranges from one to three (Figure 3). All amphibian/reptilian taxa show conspicuous within-species genetic divergence — that is, species have at least one primary phylogeographic break, and three genetic clusters are apparent in two species (Figure 3). However, populations of amphibian/reptile taxa were generally less structured than mammals, which are characterized my more monophyletic clades with the divisions coinciding with the rivers. Among the bird taxa, six of the ten species did not show population structure associated with rivers (at least based on consideration of a corresponding genealogical structure with the geographic position of rivers), whereas four species exhibited one phylogeographic division, and two species exhibited no geographic structure (i.e., individuals from different mitochondrial clades overlap geographically).

**Figure 3.**
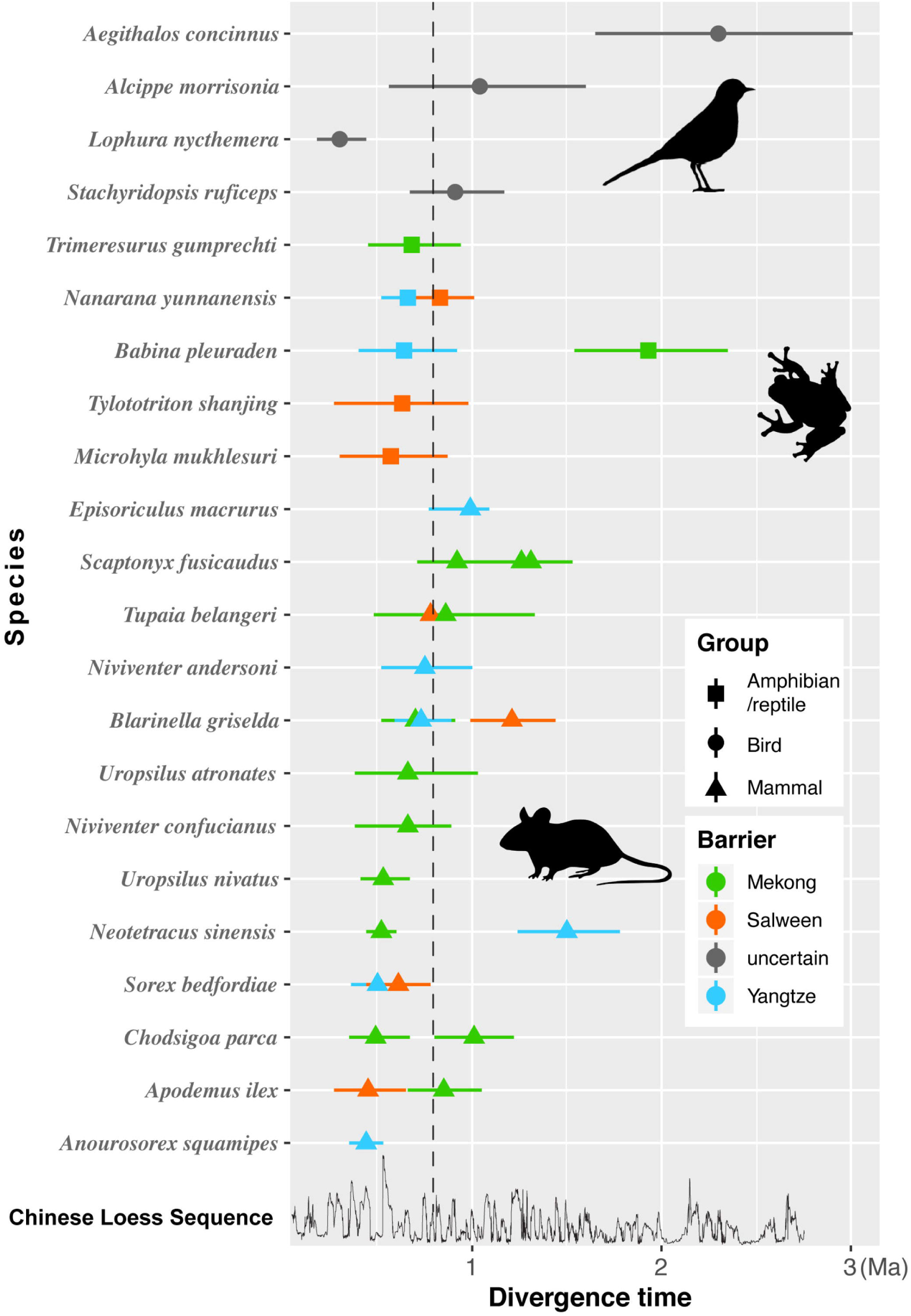
Divergence times (with mean and 95% credible intervals) of 33 phylogeographic clades cross riverine barriers; branch lengths estimated in BPP were divided by the mutation rate to express time in years. Divergence times of amphibians and reptiles, birds, and mammals, are represented by different shapes, whereas divergences associated with the different rivers are shown in differing colors. The dashed line marks the boundary of Early and Mid-Pleistocene; the bottom plot shows the climatic cycles of the Chinese Loess Sequence, where peaks represent glaciations.

With regards to phylogeographic structure associated with specific rivers, the Mekong river contributes more than the other two in shaping the phylogeographic diversity of the MSWC vertebrate community studied here (Figure 3). Specifically, 14 phylogeographic divisions are associated with the Mekong (12 of which are in mammals), 7 with the Salween (4 in mammals), and 8 with the Yangtze (6 in mammals). However, there is not a significant association between particular taxonomic groups and specific river barriers (paired t-test, *p* >0.05; Figure 3).

### Testing community-wide co-divergence in time

Using the estimated gene trees to define clades, in some cases, the clades correspond to proposed subspecies, or even potential species boundaries that are under debate. Given the taxonomic status of lineages defined by the clades does not have a direct effect on the time estimates of divergence events, we hereafter refer to all of them as clades, and which results in a total to 33 divergence events in our study (Figure 3).

For individual clade pair, we used two programs, BEAST (17) and BPP (18), to estimate the divergence time of individual clade pairs. Overall, divergence time estimates in BEAST had higher variance than those from BPP (Table S1), but the median values were similar (Figure S4, Supplementary file 2). The estimated divergence times of mammalian clades span from 0.44 to 1.50 Ma, compared with 0.57 and 0.83 Ma in amphibians and reptiles, with the single exception of an older split at 1.93 Ma in the Yunnan pond frog (*Babina pleuraden*; Figure 3). In contrast, the divergence times in birds were much more varied, ranging from 0.30 Ma to 2.30 Ma (Figure 3; Table S1). From the perspective of each of the river barriers, the timing of divergence events largely overlap in range, but clearly do not support a single divergence event for all species in the community (Salween: 0.45–1.21 Ma; Mekong: 0.49–1.93 Ma; Yangtze: 0.44–1.50 Ma; Figure 3).

To test co-divergence across species, we chose the FBC method implemented in ecoevolity (15) instead of other ABC methods based on summary statistics. Relying on insufficient summary statistics inevitably discards some information in the data, and the chosen summary statistics might be insufficient (19). FBC takes full advantage of sequence information by calculating the likelihood of the population divergence model directly from genetic data while analytically integrating all possible genealogical and mutational histories. Compared to hABC (20), this method is less sensitive to priors and less biased towards co-divergence events (15). In addition, simulation using empirically derived parameters showed that the FBC outperforms hABC with single-locus data (Jamie Oaks in prep.), and therefore we focus exclusively on the results based on FBC analyses.

We first ran FBC with all the taxa pairs multiple times with different priors on the concentration parameter of the Dirichlet process. Because of the lack of clear geographic barrier, we excluded the bird species from this comparative analysis, leaving twenty-nine mammalian, amphibian, and reptilian clade pairs. There is no posterior support for total synchronous divergence (Bayes factor <1), suggesting that the diversification at the MSWC is a complicated history. For the number of co-divergent event, 14–19 co-divergence events have highest posterior support but with considerable uncertainty — the posterior distribution is very flat and no posterior probability is larger than 0.18 (i.e., the posterior distribution is very flat; Table S3, Supplementary file 1), so the estimated number should be treated with caution.

Given this lack of temporal synchrony when considering the entire dataset, we conducted separate FBC runs for each river to examine whether the divergence events at each river were also asynchronous. The results clearly show support for limited numbers of co-divergence events (Table 1 and Figure 4). This includes: three supported co-divergence events for the Yangtze river (PP = 0.55, 2lnBF = 18.28; Figure 4), and two co-divergence events for the Mekong (PP = 0.52, 2lnBF > 27.82) and Salween (PP = 0.91, 2lnBF = 23.55) rivers (Figure 4), although there is no posterior support for synchronous divergence for any given river (Bayes factor <1; Figure 4). For any given river, we note that all the divergence events are relatively recent, but tend to include a more recent and relatively older divergence. For example, divergence events around 0.5Ma for the Salween and Yangtze, and 0.85Ma for the Mekong river, are coupled with a relatively more recent diversification pulses between 0.1–0.2 Ma for each river. This explains why the overall correlation between the temporal and spatial scale is not significant. For example, despite sharing the same geographic barrier (i.e., river), the posterior distribution of estimated divergence times actually do not have more overlap compared with the posterior of divergence times for pairs across different river barriers. For example, the mean overlap of the posterior for divergence times estimates between two clade pairs at the same river is 12%, 18% and 7% for the Mekong, Salween, and Yangtze, respectively, compared with an average overlap of 18% in the posterior of divergence times across different rivers (Figure S4 *vs.* Table S3). That is, two species sharing a geographic barrier are not more likely to have similar divergence time than those from different geographic barriers. However, the FBC analyses per river clearly show that species in the MSWC community do not have completely idiosyncratic histories (Figure 4).

**Table 1.**
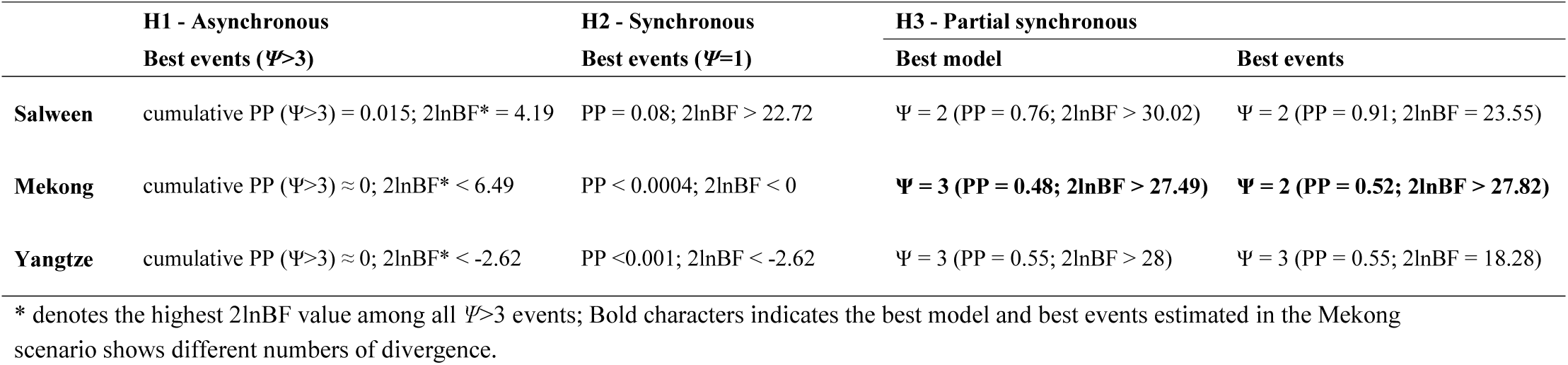
FBC results for each of the three river barriers

|  | **H1 - Asynchronous** | **H2 - Synchronous** | **H3 - Partial synchronous** |  |
| --- | --- | --- | --- | --- |
|  | **Best events ($\Psi > 3$)** | **Best events ($\Psi = 1$)** | **Best model** | **Best events** |
| **Salween** | cumulative PP ($\Psi > 3$) = 0.015; $2\ln BF^* = 4.19$ | PP = 0.08; $2\ln BF > 22.72$ | $\Psi = 2$ (PP = 0.76; $2\ln BF > 30.02$) | $\Psi = 2$ (PP = 0.91; $2\ln BF = 23.55$) |
| **Mekong** | cumulative PP ($\Psi > 3$) $\approx 0$; $2\ln BF^* < 6.49$ | PP < 0.0004; $2\ln BF < 0$ | **$\Psi = 3$ (PP = 0.48; $2\ln BF > 27.49$)** | **$\Psi = 2$ (PP = 0.52; $2\ln BF > 27.82$)** |
| **Yangtze** | cumulative PP ($\Psi > 3$) $\approx 0$; $2\ln BF^* < -2.62$ | PP < 0.001; $2\ln BF < -2.62$ | $\Psi = 3$ (PP = 0.55; $2\ln BF > 28$) | $\Psi = 3$ (PP = 0.55; $2\ln BF = 18.28$) |

\* denotes the highest $2\ln BF$ value among all $\Psi > 3$ events; Bold characters indicates the best model and best events estimated in the Mekong scenario shows different numbers of divergence.

**Figure 4.**
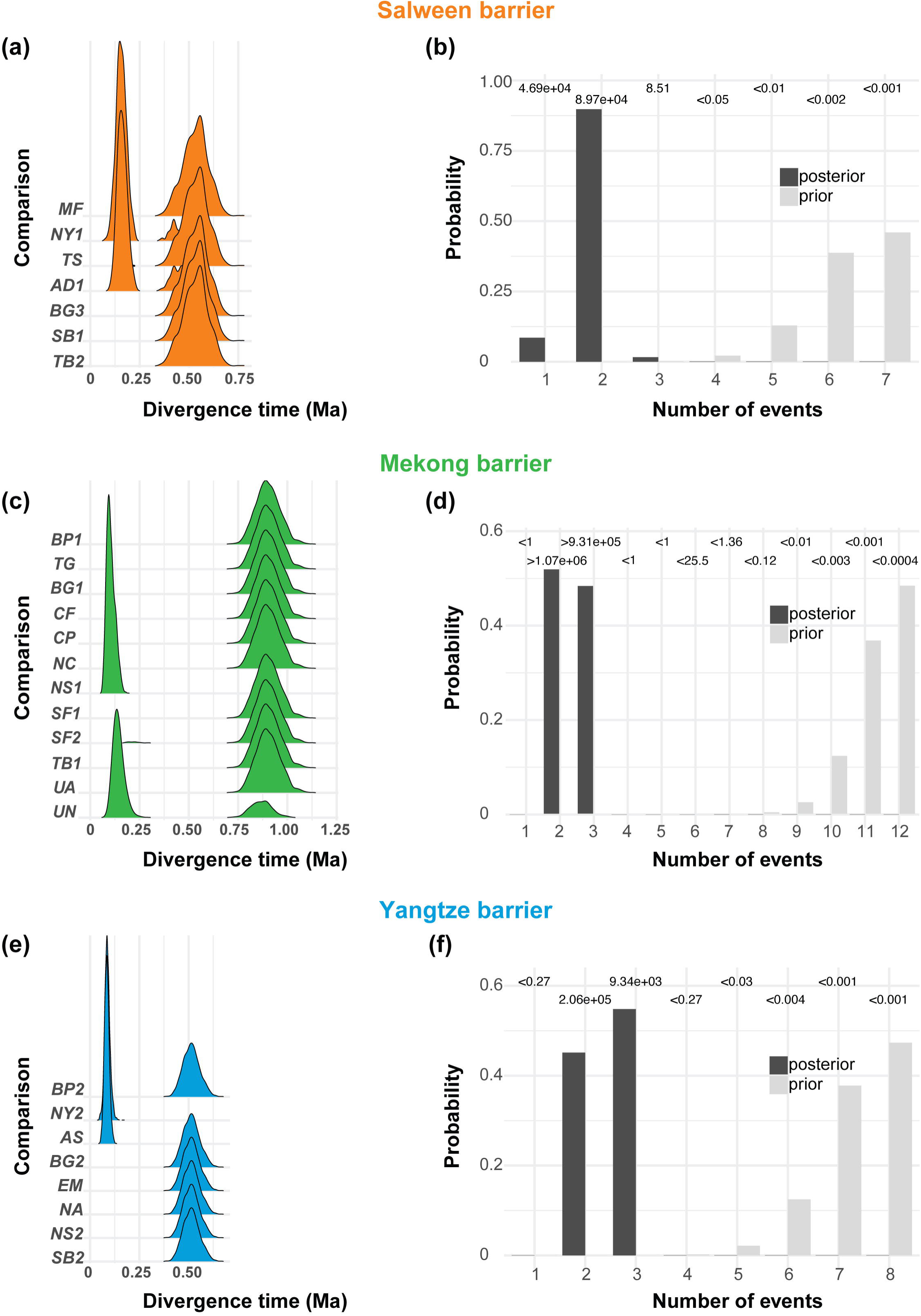
Testing phylogeographic synchronization using the full-likelihood Bayesian computation method at each river. (a), (c), (e) shows the approximate marginal posterior densities of divergence time for each clade pair. (b), (d), (f) shows the prior (light bars) and the posterior (dark bars) probabilities of the numbers of co-divergence events. The numbers above each bar denote the Bayes factor.

## Discussion

Despite more than two hundred molecular phylogenetic/phylogeographic studies to study the evolution of biodiversity in the MSWC over the past two decades (21), our study is the first to assess community-wide shared phylogeographic patterns in MSWC terrestrial animals, testing statistically whether there are shared evolutionary histories in the community. Our results reveal concordant spatial genetic structure supporting the hypothesized role of rivers as barriers structuring communities in the MSWC (Fig. S1–S3). However, this common structuring was not shared with birds in the MSWC (Table S2). Moreover, tests of temporal synchrony showed evidence of limited co-divergence (hypothesis H3 in Figure 2) — specifically, each river supported a model with multiple pulses of co-divergence events, all of which, however, occurred during the mid to late Pleistocene (i.e., 67% of the divergence events occurred between 0.5–1.0 Ma; Figure 3). This pattern is consistent with the Pleistocene “species pump” effect, where cycles of interconnections and isolation associated with climatic cycles promote divergence. Given the dozens of glacial cycles that took place in this era, the co-divergence of taxa for specific periods suggest the species pump was more pronounced during certain cycles (e.g., S1, S2, S5, S7 by Chinese Loess Sequence; MIS6, MIS8, MIS12, MIS22 by Marine Isotope Stages). Moreover, the persistence of older divergence events indicate that the permeability of river barriers varied both geographically and across taxa (Figure 3) (22). Below we discuss the implications of what this spatially concordant, but temporal dissonance, implies about diversification in the MSWC.

### Diversification in MSWC as a result of geographic features and past climate event

Climate oscillations in Pleistocene have long been known as an effective speciation mechanism in mountainous regions (23, 24). Climate induced species range shifts could lead to isolation (e.g., isolation between different refugia) or gene flow (e.g., sky island dwellers moved down to the valley forming one population) (25). Hence it is not surprising that phylogeographic studies on organisms in other mountains system (e.g., the Alps in Europe and Madrean Sky Island mountain ranges in North America) usually reveal genealogical splits possibly representing different refugia populations, and the estimated divergent time often fall within the Pleistocene (26-28). In the MSWC, any climate-induced distributional shifts (and associated refugial areas) are structured by the rivers (Figure 3) (i.e., there is a general spatial correspondence between the genealogical splits and the distribution of clades on either side of the river barriers). The primary exception to the role of rivers in MSWC structuring divergence was observed in birds, which showed less spatial concordance of phylogeographic structure and/or very little evidence of geographic structuring of genetic variation (i.e., lack of genealogical splits; Table S2).

Despite the spatial concordance of divergence patterns for vertebrates in the MSWC, the lack of temporal synchrony suggests the effectiveness of river barriers varies. We propose that a potential explanation lies in the mountains themselves. Specifically, the mountains have a north-south orientation and all have altitudinal zonation spanning from tundra to tropical forests. Both features are very rare in the world and has no doubt contributed to substantial stability of ecosystems in the MSWC. For example, for moderate glacial cycles, the majority of the MSWC species only have to migrate vertically or slightly southward to track climate change (29), given that the species richness concentrates at the mid□elevations (30-32) (see Fig. 5). Even if these animals reach the lower altitudes of the mountains, the rivers can still prevent gene flow between different populations. That is, the environmental gradients in the MSWC act as a buffer against extinction in the face of climate change and the rivers as barriers to secondary contact (9). Only during dramatic glaciations in which species’ distributions would have been displaced to the lowest altitudes, as well as latitudinally where the southern mountains are shorter (Figure 5), were there opportunities for connectivity across rivers (i.e., the river barriers become permeable). As a result, the divergence time between contemporary populations clustered with the severe glaciation events, resulting is some degree of synchrony (Figure 4). This contrasts with examples from the Amazon, another biodiversity hotspot, where rivers act as barriers (33), but divergence is asynchronous (34). We suggest the extremely rugged terrain and rivers made the impact of glaciations on species community more punctuated in MSWC, generating some clusters of divergence times (i.e., co-divergence pulses).

**Figure 5.**
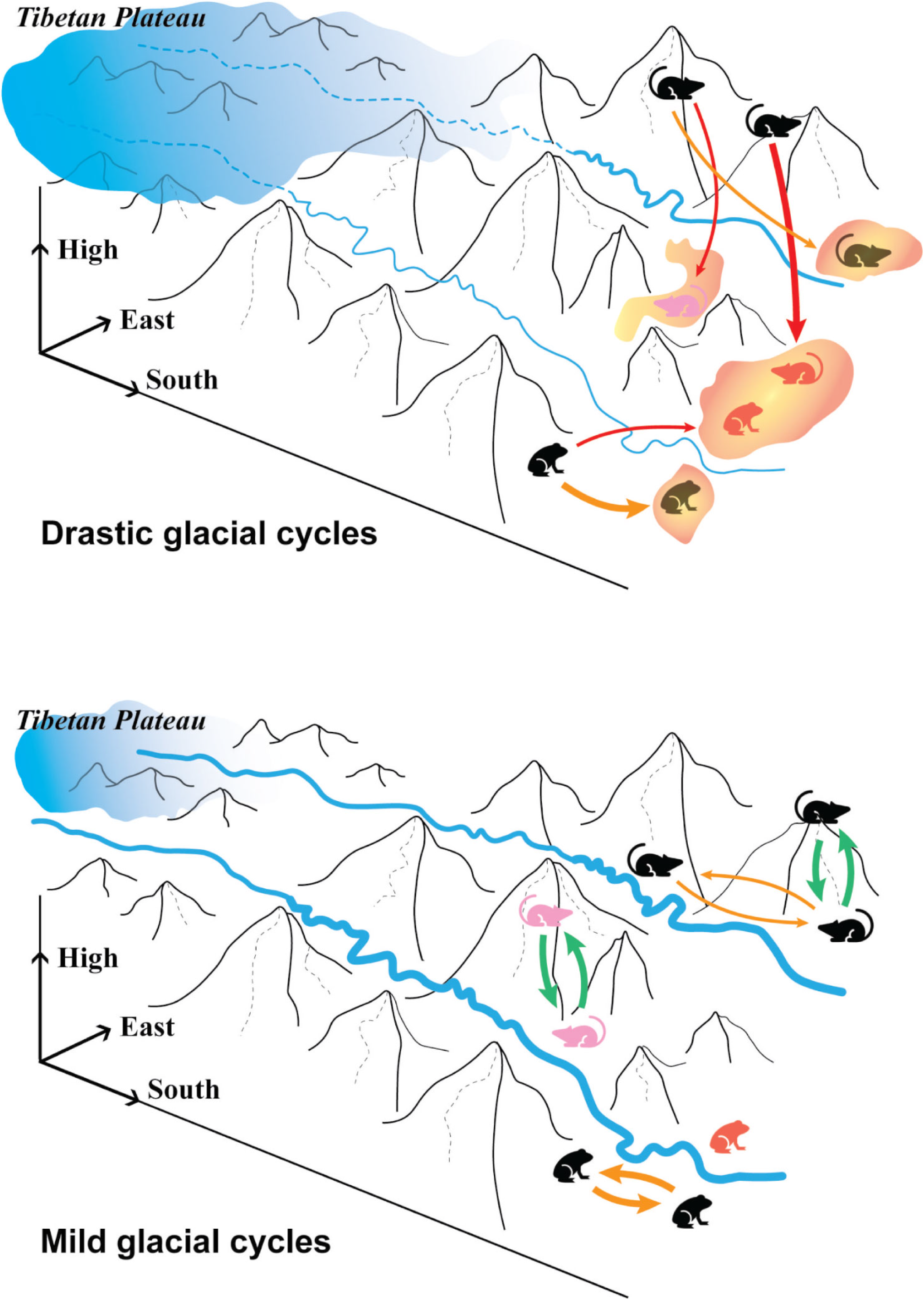
Differences in the permeability of river barriers as a function of the magnitude of climate-induced distributional shifts associated with the Pleistocene glacial cycles in the MSWC. During drastic glacial cycles, phylogeographic divergence of montane species included dispersal across river barriers (marked by the red arrows) as species tracked their climate niche altitudinally, but also latitudinally where the mountains are shorter (i.e. refugia, marked with red orange patches). This contrasts with mild glacial cycles, the environmental pressure is not harsh enough to promote cross-river dispersal. Instead, species could track climatic niche additionally using elevational gradients (green arrows).

### The importance of studying recent diversification in MSWC

Understanding the mechanism of recent diversifications is indispensable for understanding how biodiversity was generated and maintained in the MSWC. Divergence events in the MSWC encompass a broad time range, even for phylogeographic divisions within species. For example, phylogeographic divergences in subnival plants across the Mekong–Salween Divide — a classic biogeographic boundary, also known as the Ward Line — varied from late Pleistocene (0.37– 0.48 Ma) (35) to the Pliocene (4.49 Ma) (36). Mosbrugger et al. (9) recently reviewed the Cenozoic evolution of geobiodiversity in the general Tibeto-Himalayan region and pointed out the three peaks of diversification: the first flourishing of modern families and genera during the mid□Miocene (20–15Ma), the second during the Miocene-Pliocene boundary, and the third, the focus of our study, at the early and mid-Pleistocene (2.0Ma–150ka).

However, the mechanisms behind the most recent peak of diversification are often overlooked. One main reason is that MSWC is adjunct to Tibeto-Himalayan region, and the majority of the phylogeographic studies in this region focus on finding the footprint of the orogenesis process on biodiversity. Many studies reported young divergence events (2.0 Ma–150ka), but were “mired” by this “uplift story”, and falsely ascribed them to orogenesis processes. Only recently, careful reviews of geographic evidences questioned the often-cited literature on the third uplift phase of Tibetan Plateau (15–0.5 Ma) (21), and the current geographic consensus is that the orogeny in this region finished long before the Pleistocene. This leaves very few studies recognizing and investigating the effect of Pleistocene climate change on species diversity in MSWC.

Studying recent divergence (i.e. phylogeographic processes) in MSWC is also difficult, yet crucial for conservation. Many taxa in this region have taxonomic ambiguity. In fact, more than half of our studied species are cryptic, and many are still in placed in species complexes. Given the MSWC is a crucial region for conservation—over 30% of plant species (∼12,000 species), and about 50% vertebrates (∼1,100 species) of China live here (according to the Catalogue of Life China 2019 Annual Checklist), including many endangered species. Characterizing these cryptic diversities—how they are distributed spatially and how climate change might affect them— is important for comprehensive conservation planning in the MSWC (37, 38).

### Future directions

Our study provided insights on the MSWC’s often-neglected recent diversification process by studying both spatial and temporal aspects of concordance across the community. One drawback due to the availability of published data is that we only used mitochondrial loci in the analysis, which limits the accuracy of divergence time estimates. By analyzing a “community” of single-locus dataset, we gain some statistical power, and by applying the recently developed FBC method (as opposed to relying on approximation approaches), we are better able to extract information from the genealogical histories across taxa (15). Nevertheless, and with the expansion of genomic data in phylogeographic studies (e.g., RAD-seq) (39), the robustness of our findings needs to be confirmed (40).

Future studies should also consider expanding the taxon sampling to evaluate the extent to which temporal dissonance is nevertheless characterized by some co-divergence pulses during different glacial periods. More taxa also would make it possible to explore the connection between biological traits and divergence pulses (14, 22, 41). Such analyses would be especially interesting for addressing why the permeability of river barriers varies. That is, with the signature of older and more recent divergence pulses (Fig 4), it is an outstanding question why some taxa (but not all) did not disperse during the recent connections that some taxa obviously used to disperse across the river (i.e., the timing of divergence dates to the recent past).

## Conclusion

With complex terrain, mountains harbor a substantial proportion of the world’s species. This disproportionately high species richness, as well as the mechanisms of evolutionary diversification, have long intrigued biologists (42, 43). With the help of comparative phylogeographic methods, our study explored the fascinating phylogeographic diversity harbored in the MSWC and identified that spatial concordance is not associated with temporal synchrony. Instead, multiple pulses of co-divergence, which coincide with the more pronounced glacial cycles suggests a complex biogeographic process akin to a Pleistocene’s species pump effect across the extremely rugged terrain of the MSWC. Future studies in the MSWC with genomic-scale data and more species sampling would enhance our understanding of the accumulation of mountain biodiversity evolution across space and time.

## Materials and Methods

The dataset included mitochondrial DNA (mtDNA) data from 24 species collected from GenBank (see Table S1 for the Genbank no. and original studies). The present study also sequenced the Cytb gene in six mammal species (Genbank no. XXX-XXX).

### Matrilineal phylogeographic structure and divergence time estimation for individual species

We analyze the phylogeographic breaks for each species in two steps. First, we estimated a gene tree using BEAST v.2 (16). If the original paper did not specify, we set the prior distribution of substitution rate with a mean of 0.018 substitutions/site/Ma for mammals (44), 0.0105 substitutions/site/Ma for birds and reptiles (45) and 0.010 substitutions/site/Ma for amphibians (46), and a standard deviation of 0.005. All trees were time-calibrated using an uncorrelated relaxed clock model. In each species, the analysis was run 20 million generations with a sampling frequency of 2000, discarding the first 20% as burn-in. Further, phylogeographic clades were cross-checked using the GMYC method implemented in the “splits” R package (17). The ultrametric tree from BEAST was used as the input for GMYC estimation, both single- and multiple-threshold for delimiting were applied, and we only considered the more conservative delimitation. In addition to BEAST, BPP v3.4 (18) was also used to estimate the divergence time. We set the gamma prior on the population size parameter *θ* according to empirically estimated Watterson’s *θ* summary statistic. Because of limited information for the divergence time *τ*, we determined its gamma prior by running preliminary runs in BPP to compare a deep divergence scenario with *τ* ∼ G (2, 200) and a shallow divergence scenario with *τ*∼G (2, 2000). The scenario with converged MCMC chains was chosen for each run. BPP analyses were run for 100,000 generations following a burn-in of 10,000 with a sampling frequency of 5. The convergence of *τ* and *q* parameters were examined in Tracer 1.6 (47), and estimated *τ* was converted to absolute time following Burgess & Yang (48) with the mean mutation rate of each species.

### Detecting community-wide phylogeographic concordance

On our empirical data, we ran three independent MCMC chains for each river-specific analysis as well as the all-in-one analysis to verify convergence among the MCMC chains; 12 FBC runs were conducted in total. For each analysis, we used the following settings: (i) we selected the value of the Dirichlet process concentration prior to place approximately 50% of the prior probability on the maximum number of divergence events using dpprobs program of the ecoevolity package (15); (ii) a gamma distribution (shape = 2.0, scale = 0.005) as the prior for population size, which is expected to be broad enough to cover the population sizes of studied species; (iii) a global set of mutation rate (μ) as 0.02/site/Ma and a scaler for individual clade pairs was applied via their mutation rate relative to the global setting. All the other parameters were set as default (i.e. exponential distribution of event time prior with a rate of 100, auto optimization of operator setting with delay value of 1000). We used 150,000 generations for MCMC with a sampling frequency of 50, the convergence of MCMC chains were checked using Tracer (47).

For the FBC analysis with all the clade pair, we expected an over-estimated number of co-divergences. Applying a temporal “buffer” between pulses of co-divergences would filter the noise when large numbers of clade-pairs are submitted to ABC computation (49), but is not available with the current version of FBC. Hence, to investigated whether co-divergence on the temporal scale corresponds to co-divergence on the spatial scale, we extracted the posterior distribution of time estimates for each clade pair and calculated overlaps between the distributions. This gives a measure of how likely two clade-pairs have similar divergence time, more quantitative then directly using the FBC estimated co-divergence events. The FBC analyses for each riverine barrier were conducted with similar setting.

## Acknowledgements

The authors acknowledge Knowles lab for constructive discussions and comments on this study. In particular, we are grateful for Dr. Luciana Moreira and Dr. Andréa Thomaz for their help on the analysis of comparative phylogeographic model, and Dr. Richard Holder for the help in illustration of the results. We also express thanks to the high-performance computing resources and other facilities provided by the Department of Ecology and Evolutionary Biology, University of Michigan, as well as Dr. Kai He for his suggestions in the design of the research.

