## Supplementary file 2 for "Neglected Quaternary legacy on biodiversity in the Mountains of Southwest China"

**Neglected Quaternary legacy of biodiversity for the Mountains of Southwest China**

Tao Wan, Huateng Huang, Jamie Oaks, Xuelong Jiang^#^, L. Lacey Knowles

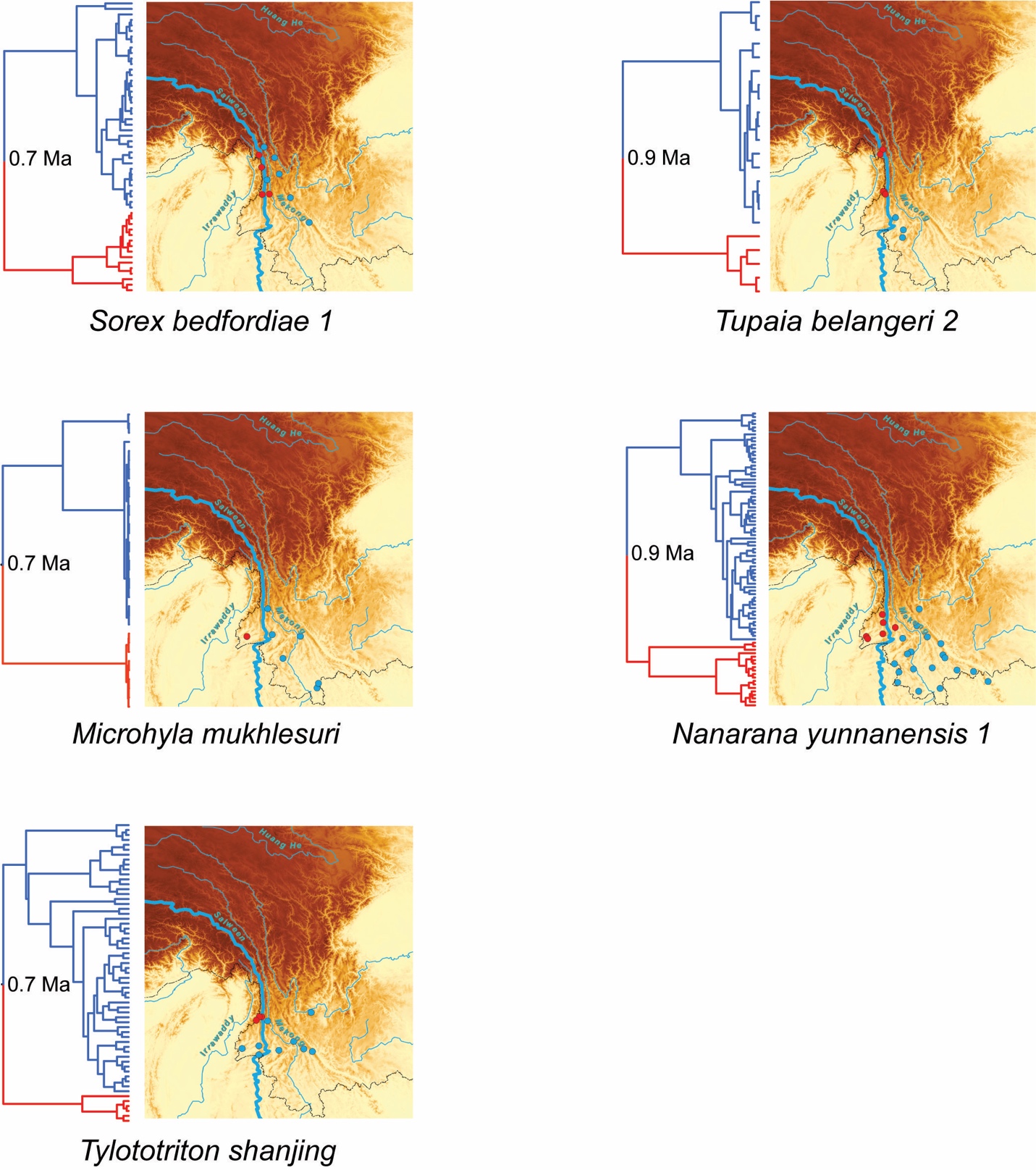

**Figure S1**. Time-calibrated tree of the taxon that under the isolation of Salween River, indicating crown age, phylogenetic relationship, and location of samples and geographic break.

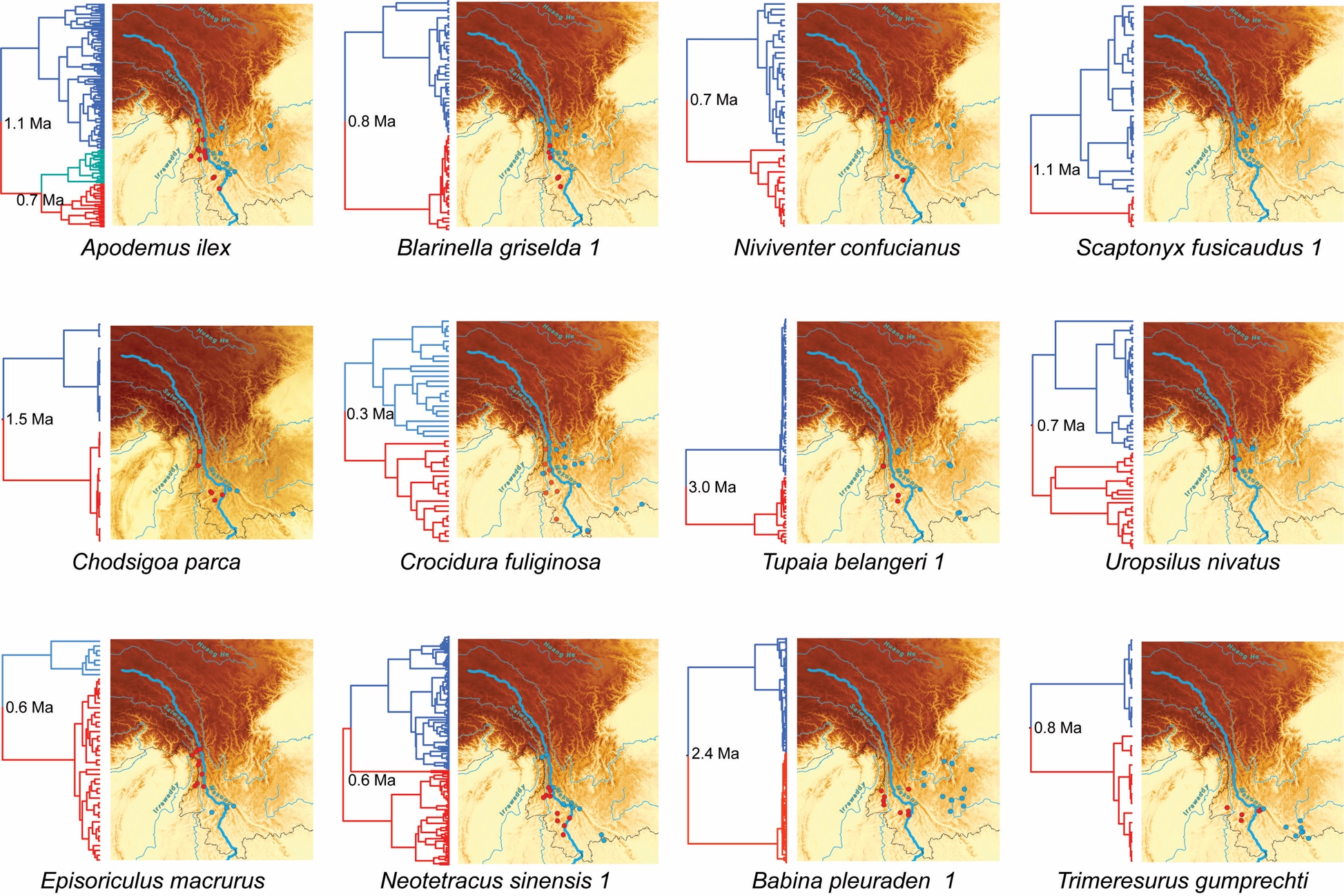

**Figure S2**. Time-calibrated tree of the taxon that under the isolation of Mekong River, indicating crown age, phylogenetic relationship, and location of samples and geographic break.

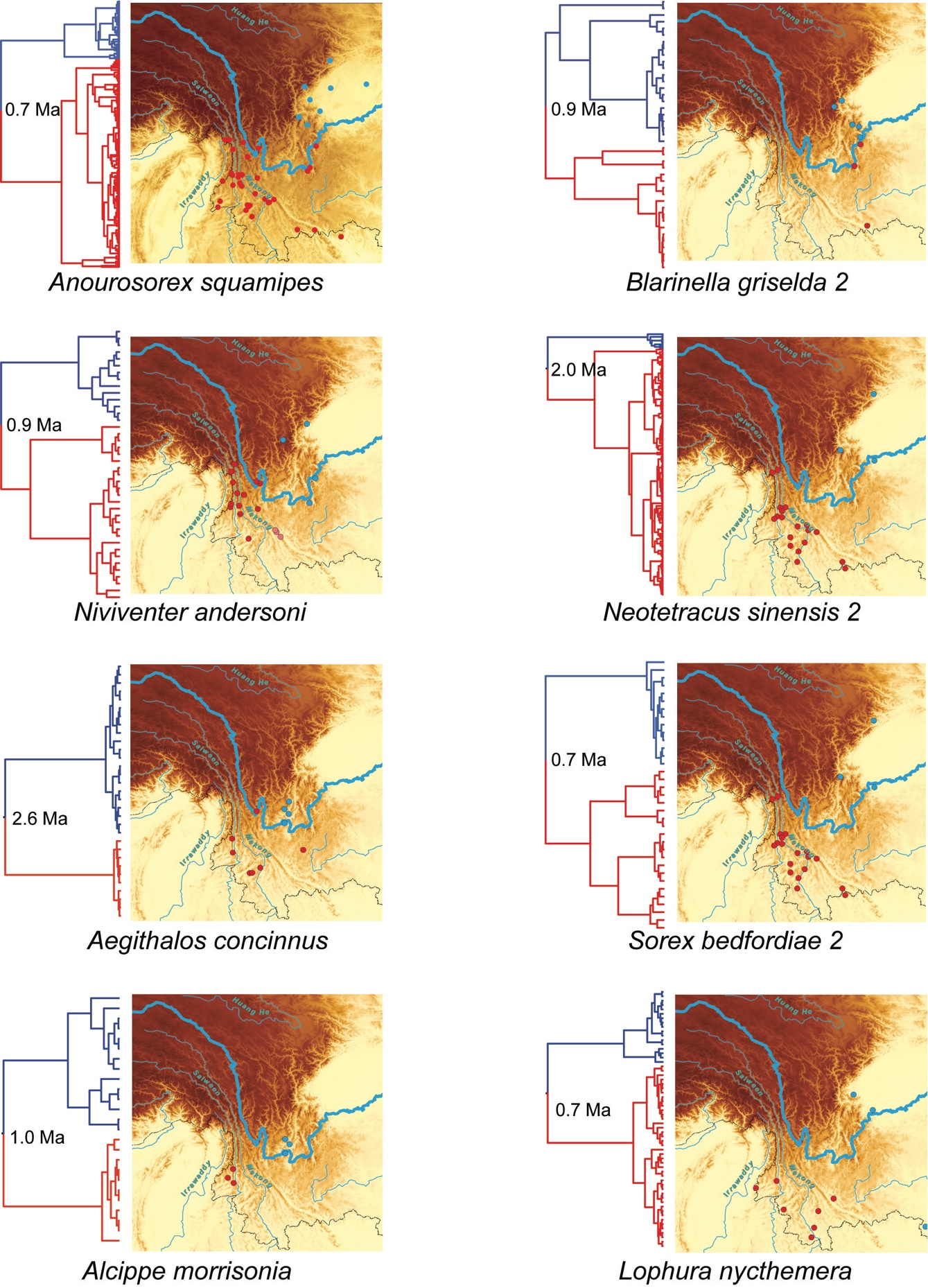

**Figure S3**. Time-calibrated tree of the taxon that under the isolation of Yangtze River, indicating crown age, phylogenetic relationship, and location of samples and geographic break.

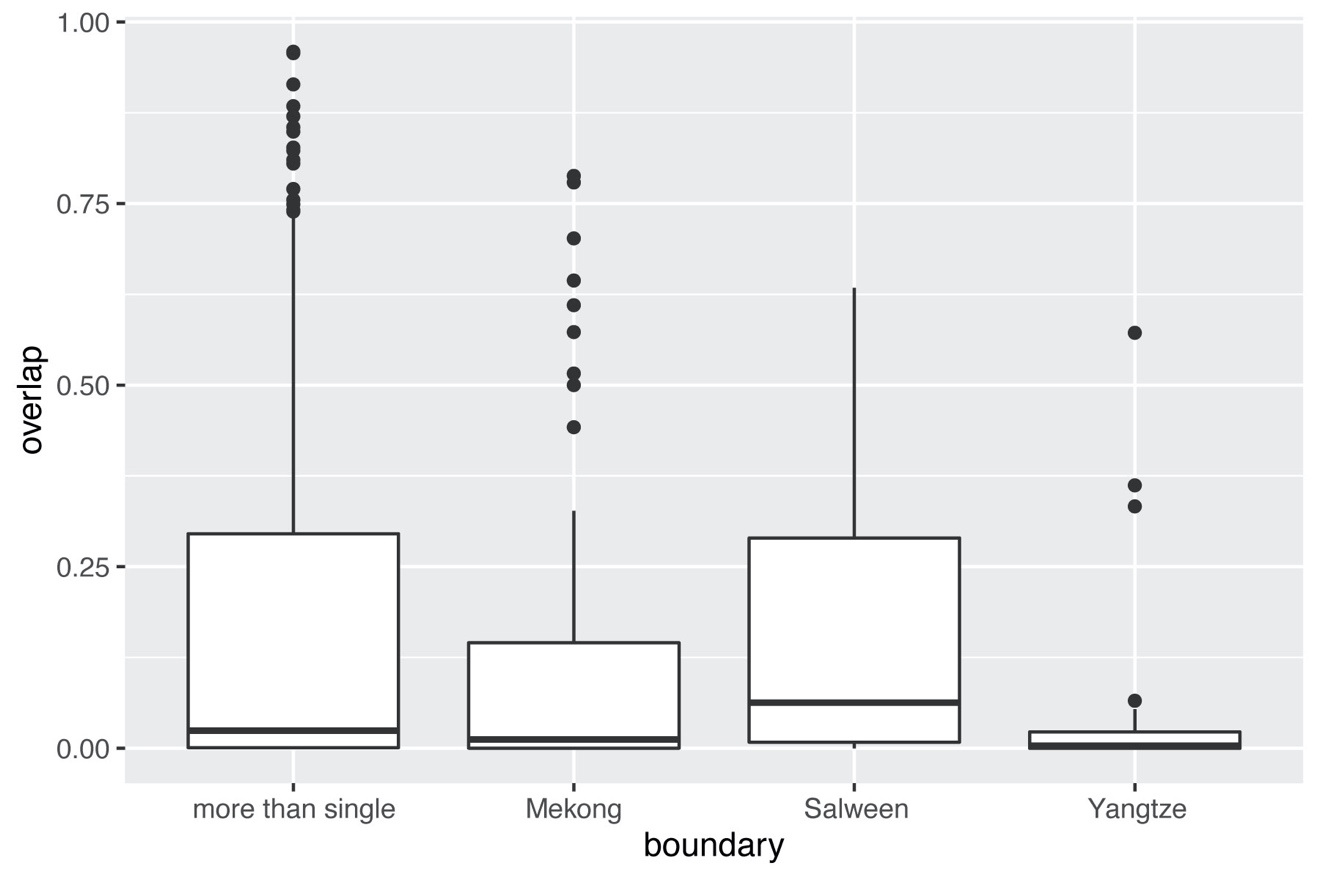

**Figure S4**. Percentage of divergence time overlap between clade-pairs with share and non-share single riverine barrier.
